# A novel methodology for assessing reinforcement choice in primates under human care

**DOI:** 10.64898/2026.10.30.763045

**Authors:** Maria Isabel Leiva, Rowland K. Griffin, Todd R. Lewis

## Abstract

Positive Reinforcement Training is a technique commonly employed in zoological collections to elicit a desired behaviour from an animal under human care. Typically, a food item is offered as a reward for performing that behaviour. Three non-human primate species (*Mandrillus sphinx, Pan troglodytes*, and *Hylobates lar*) were offered four different foods (banana, grape, apple, and sweet potato) in a series of choice experiments. Latent variable models were employed to investigate which foods were the best selection to use as reinforcement items. Results show that food preference varies between the three primate species. *Pan troglodytes* and *H. lar* showed strong preferences for grape and banana, whereas *M. sphinx* showed a very strong preference for sweet potato. This study indicates that foraging ecology might play an important part in fruit selection and preference in primates. The findings highlight that understanding a species’ natural behaviour and dietary preferences is key to informing reinforcement and training decisions.

## 1. Introduction

Animal training can be defined as intentionally modifying an individuals’ behaviour, whilst having an understanding of behaviour analysis and the successful application of this philosophy (Westlund 2015; Heidenreich et al. 2023). The goal of animal training is to modify the behaviour while keeping animal welfare in mind. One commonly used technique for training animals under human care in zoological collections is Positive Reinforcement Training (PRT) (Savastano et al. 2003; Schapiro et al. 2003; Westlund 2015). This enables zookeepers and veterinarians to perform routine health checks, administer medication, and enrich the lives of these animals (Savastano et al. 2003).

The use of PRT techniques in non-human primates can be particularly important as these animals are often dangerous to work with when in close proximity. As such, positive reinforcement can significantly aid in the safe management and welfare of these animals in zoological collections. Whilst there is breadth of research into the efficacy and benefits of PRT techniques in primates, there remains a paucity of information regarding food preferences during these interactions.

Traditionally, primates housed in zoological collections have been fed high fruit diets mixed with vegetables, insects, and vertebrate prey (Plowman 2013; Britt et al. 2015). As a result, these diets were often high in sugar and low in protein and fibre (Schwitzer et al. 2008).

Over the last decade or so, it has become apparent that these types of foods are unsuitable for the nutritional and energetic requirements of most primate species and that such diets have led to a multitude of health and behavioural problems (Plowman 2013; Britt et al. 2015; Viallard et al. 2023). This realization has led to a movement away from high sugar, low protein/fibre diets, with many nutritional departments designing fruit-free diets that balance energetic and nutritional needs of primates in a more naturalistic way (Plowman 2013).

In 2023, the primate collection in Parque Zoológico Nacional la Aurora (PZNA) was transitioned from a traditional fruit-based diet to a fruit-free diet. The philosophy behind the change was to provide a more naturally balanced nutritional range that reflects their diets in the wild. The diet of primates at PZNA now consist of 95% non-fruit components, with the final 5% consisting of fruit that is used for positive reinforcement. In this study we assessed which fruits were the best selection to use as reinforcement items by employing a choice experiment to test the hypothesis that fruit preference will differ depending on the species of primate.

## 2. Method

Parque Zoológico Nacional la Aurora is located in Guatemala City, Guatemala and the collection includes 13 primate species. During this study we focused on three species, mandrill (*Mandrillus sphinx*), chimpanzee (*Pan troglodytes*), and white-handed gibbon (*Hylobates lar*). These species were selected due to: a) their small group sizes which enabled choice selection to be recorded effectively; and b) their differing natural history traits: mandrills are a primarily terrestrial species, chimpanzees semi-arboreal, and gibbons arboreal. The group sizes for each species were as follows: mandrill (2 males), chimpanzee (2 males and 3 females), and gibbon (2 males and 1 female). Four fruits (grape, banana, apple, and sweet potato) were used in the assessment. All animals studied were housed at PZNA and remained in their usual enclosures with normal husbandry throughout the study.

Choice events consisted of 10 replications of choice with a combination of two of the four fruits. Every combination of fruit was offered as a choice event to each species twice. Thus, a total of 36 choice events (12 per species) were conducted within 20 days. Each replication of choice was offered to a specific individual within the group and the first choice of fruit was recorded each time. The choice events lasted an average of 15 minutes each, summing up to three hours of observation per species. Choice events were conducted at random times that were not associated with regular feeding times over a period of nine weeks, starting from April 5 until June 7 2024. The same animal trainer conducted the choice events to avoid observer bias.

For analysis, Generalized Linear Latent Variable Models (GLLVM) were performed in R package “gllvm” (Niku et al. 2019a; R Core Team 2021). GLLVM is a model-based multivariate method that has been successfully applied to previous animal behaviour studies (Rice et al. 2022; Jenner and Lewis 2024; Hughes et al. 2025). GLLVM extend basic generalized linear models on multivariate data using a factor analytic approach by incorporating latent variables to combine values with factor loadings that model correlation between responses. These latent variables have a natural interpretation as ordination axes and can predict new values, control for known variables, and assist model selection (Hui et al. 2015; 2017). Primate behaviour multivariate data were constructed as a matrix with n rows (fruit choice) and m columns of behaviour frequency response during choice events.

The GLLVM regressed the mean species frequency of behaviour against a matrix of predictor factors of food type. Models used negative binomial, Poisson, and zero-inflated Poisson (ZIP) families via package TMB, with variational approximation or Laplace distributions (Kristensen et al. 2016). Dunn-Smyth residuals and Quantile plots were used to inspect model fit. Model fit was further assessed by AIC/BIC, and a for-loop iteration used to select appropriate numbers of latent variables, in this case N= *2* for all models. Models used a best of five run routines with the highest log-likelihood model selected (Niku et al. 2019b). The getResidualCor function was used with package corrplot (Wei and Simko 2017).

The final model was selected based on residual fit, AIC/BIC, moderation of correlation, suitable number of latent variables, and then estimated coefficients for predictors plotted with their confidence intervals and used for interpretation. Based on model fit from the Q-Q plots and AIC/BIC scores, the ZIP model was chosen for interpretation of results. Model fit plots and tables related to analysis can be viewed in the supplementary material.

## 3. Results

The coefficient plots from the ZIP model revealed significant results in terms of fruit preference in all three species of primates tested (Figure 1). Interestingly, while there are similarities between species, there were noticeable differences in responses to different fruits. Gibbons showed a strong preference for grape, followed by banana, and no preference for apple or sweet potato. Chimpanzees showed preferences for grape, followed by banana, with no preference for apple or sweet potato. Whereas Mandrills showed a strong preference for sweet potato and weaker preferences for grape, and no preference for apple and banana (Figures 1 and 2).

**Figure 1.**
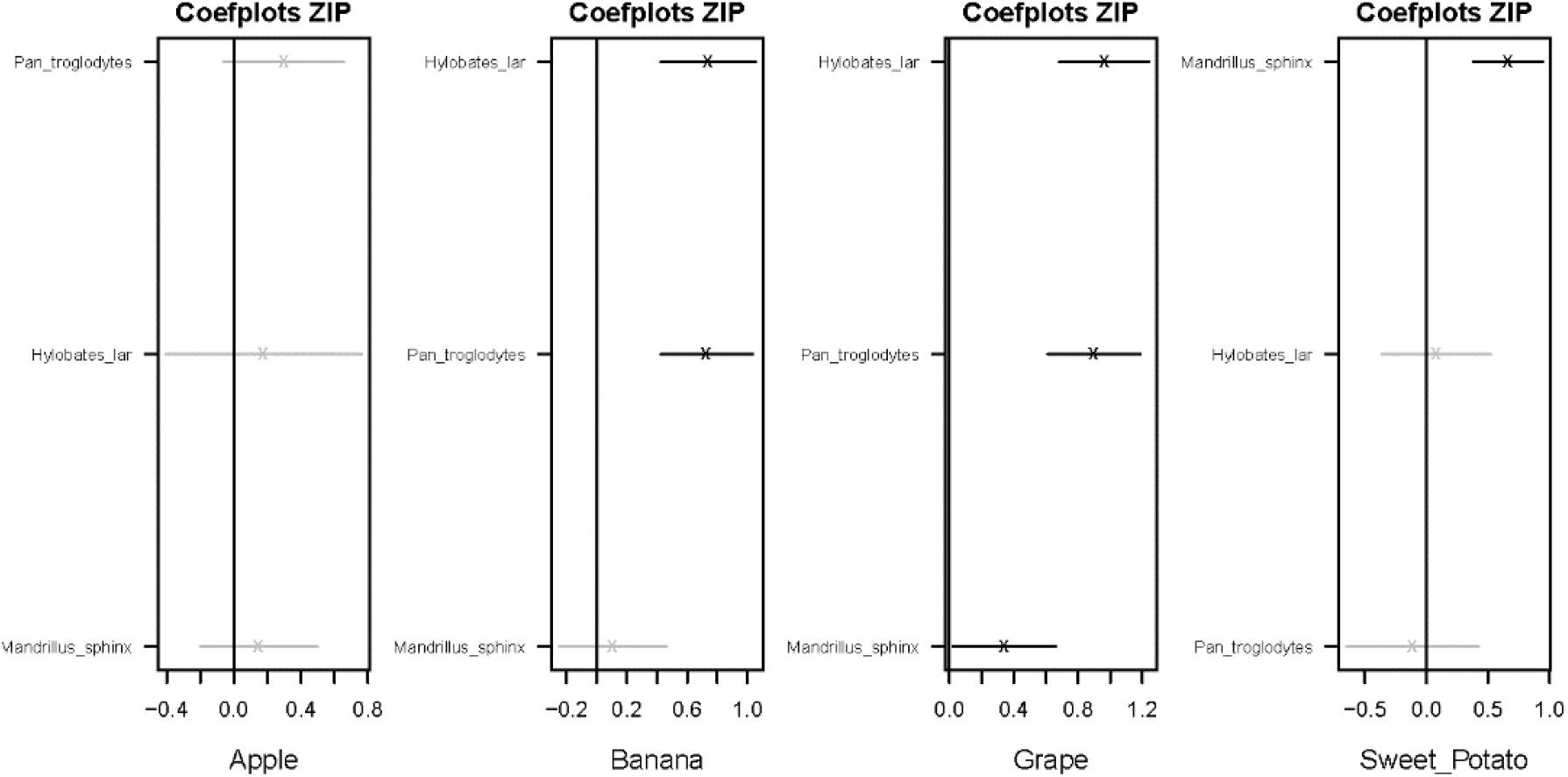
Latent variable coefficient plots with confidence intervals (horizontal lines) showing associations between primates and fruit choice. Significant coefficients are indicated in bold, and their confidence intervals do not cross zero. Coefficients that cross zero, and are therefore not significant, are indicated in grey.

**Figure 2.**
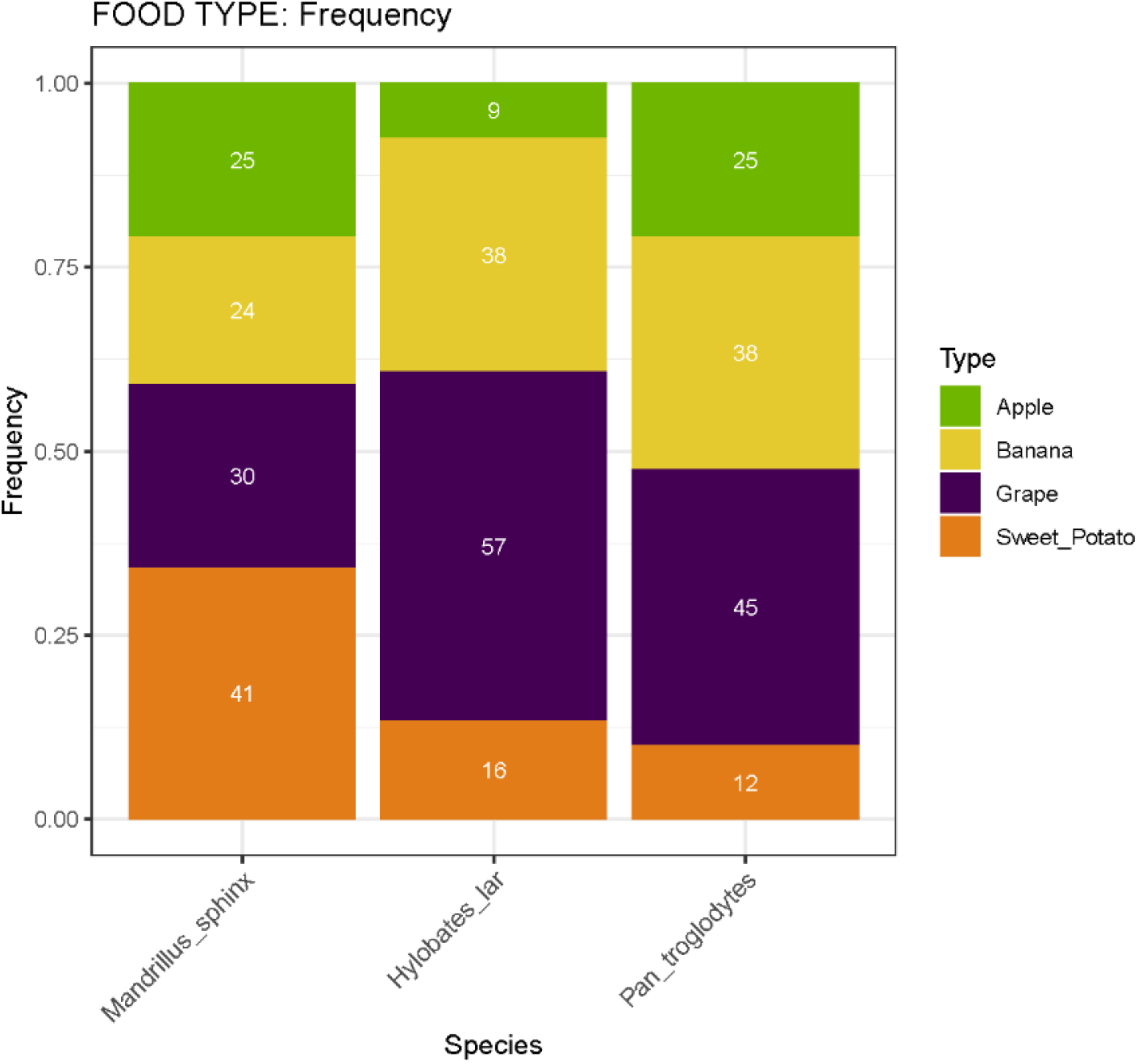
Barplot showing the frequency, represented by percentage, each fruit was chosen by three primate species. White numerals in the columns indicate the number of times each fruit was chosen by that species.

## 4. Discussion

This study shows that fruit preference varies among primate species. This is corroborated in various studies that show differences in colour recognition related to fruit colour change depending on the species of primate being tested (Hernández et al. 2021). Studies on white-handed gibbons under human care show a preference for fruit based on carbohydrate, fructose and glucose content of the respective fruit, with grape, banana and fig being preferred over other fruits (Jildmalm et al. 2008). Assessments of wild diet in chimpanzees had been shown to vary seasonally showing a strong preference for ripe fruit when it is available in their environment (Conklin-Brittain et al. 1998). More recently, studies have attempted to link the effect of seasonal dietary preferences to the quality of well-being and health in captive chimpanzees (Gerstner and Pruetz 2022). It would be interesting to investigate whether these temporal shifts in natural diet affect fruit preference response in a zoological setting.

The results of this study loosely conform with previous studies and show that at PZNA white-handed gibbons and chimpanzees exhibit preferences for grape and banana, whilst mandrills exhibit a very strong preference for sweet potato over all other fruit offered (Conklin-Brittain et al. 1998; Jildmalm et al. 2008; Nsi Akoue et al. 2017; Hongo et al. 2018). The preference for sweet potato in mandrills seen in this study may be influenced by conspecific dominance within the group, as one male in this group is highly dominant of the other. As such the second male rarely gained the opportunity to make a choice selection during this study. Such bias can be considered a limitation and we recommend adjustment of this for any future study.

This study suggests that foraging ecology might play an important part in fruit selection and preference in primates. For example, mandrills are a mainly terrestrial foraging species that spend substantial time looking for food in the leaf litter of the forest floor (Nsi Akoue et al. 2017; Hongo et al. 2018). A preference for sweet potato is in-keeping with this foraging strategy. Equally, the preferences for grapes and bananas observed in gibbons and chimpanzees which exhibit arboreal and semi-arboreal ecologies, fits this notion (Barelli et al. 2008; McLennan 2013).

Importantly this study shows that understanding a species’ natural behaviour and dietary preferences is key to informing reinforcement and training decisions, and that those preferences are not necessarily appropriate to a closely related species without careful consideration of that species’ preferences and ecology.

## 5. Author Contributions

Maria Isabel Leiva: Conceptualization, data curation, formal analysis, investigation, methodology, project administration, resources, software, validation, visualization, writing – original draft, writing – review and editing. Rowland K. Griffin: Conceptualization, formal analysis, investigation, methodology, resources, software, supervision, validation, visualization, writing – original draft, writing – review and editing. Todd R. Lewis: Formal analysis, investigation, methodology, resources, software, supervision, validation, visualization, writing – review and editing.

## 6. Acknowledgements

MIL and RKG would like to thank the Board of Directors of Asociación Guatemalteca de Historia Natural and Parque Zoológico Nacional la Aurora for their support of this project. TRL acknowledges UWE Bristol for support.

## 7. Ethical Statement

This study was a feeding trial of animals housed in a long-term managed care setting. No invasive procedures were conducted. Animal care was provided by regular qualified staff at Parque Zoológico Nacional la Aurora, Guatemala. All animals remained at the zoological facilities upon completion of this study. Parque Zoológico Nacional la Aurora is a Charity registered in Guatemala (Resolución No. 1301-90). It is licensed by the Congress of the Republic of Guatemala and its actions approved under Animal Welfare Law (Decree 5-2017).

## 9. Supplementary Material

The data and R code that support this study are available open access at: https://doi.org/10.5281/zenodo.23732758

